# Assessing the quality of *de novo* parasite genomes assembled using only Oxford Nanopore Technologies MinION data

**DOI:** 10.1101/2023.10.12.562065

**Authors:** Kaylee S. Herzog, Rachel Wu, John M. Hawdon, Peter Nejsum, Joseph R. Fauver

## Abstract

Parasitic nematode infections represent a significant burden of disease in impoverished populations. Genomic studies of parasitic nematodes have revealed novel drug and vaccine targets and provided unprecedented insights into parasite biology. A key component of these studies is the availability of high-quality reference genomes (i.e., genomes that are contiguous, complete, and accurate) that capture the biological diversity contained within species. However, relatively few genomic resources exist for parasitic nematodes and few species are represented by more than a single reference genome. Streamlined laboratory and computational workflows to generate high-quality reference genomes from individual specimens using a single data source have the potential to increase the availability of genomic data from parasitic nematodes. The Oxford Nanopore Technologies (ONT) MinION is an accessible sequencing platform capable of generating ultra-long read data ideal for assembling genomes. However, lower read-level accuracy of ONT data has previously required assemblies to be error-corrected with more accurate short-read data. In this study, we assessed the quality of *de novo* genome assemblies for three species of parasitic nematodes (*Brugia malayi, Trichuris trichiura,* and *Ancylostoma caninum*) generated using only ONT MinION data. Assemblies were benchmarked against current reference genomes and against additional assemblies that were supplemented with short-read Illumina data through polishing or hybrid assembly approaches. For each species, assemblies generated using only MinION data had similar or superior measures of contiguity, completeness, and gene content. In terms of gene composition, depending on the species, between 88.9–97.6% of complete coding sequences predicted in MinION data only assemblies were identical to those predicted in assemblies polished with Illumina data. Polishing MinION data only assemblies with Illumina data therefore improved gene-level accuracy to a degree. Furthermore, modified DNA extraction and library preparation protocols produced sufficient genomic DNA from *B. malayi* and *T. trichiura* to generate *de novo* assemblies from individual specimens.

**Data Availability Statement:** Quality-controlled MinION and Illumina data for each species are deposited in the NCBI Sequence Read Archive (SRA) under the BioProject accession ID PRJNA1074771 under BioSample accession nos. SAMN39888962 (*Brugia malayi*), SAMN39888963 (*Trichuris trichiura*) and SAMN39888964 (*Ancylostoma caninum*). Final assemblies for each species are publicly available on GenBank.

## Introduction

Parasitic nematodes represent an enormous burden of disease. It is estimated that 1.5 billion people are infected with soil-transmitted helminths, while approximately 120 million and 15 million people suffer from filariasis and onchocerciasis, respectively^1^. Helminths are responsible for nearly 12 million Disability-Adjusted Life Years (DALYs) lost annually^1^. In addition to the morbidity caused by nematodes themselves, infection also exacerbates the disease burden of other existing conditions such as malaria, HIV, and tuberculosis, and can reduce immunological response to important vaccines^2–6^. Reference genomes for parasitic nematodes have proven to be an invaluable resource for increasing our understanding of helminth biology, treatment, and control. For example, the landmark comparative genomic study by the International Helminth Genomes Consortium^7^ analyzed dozens of complete genomes representing all major helminth lineages. This extensive dataset allowed for the identification of multiple gene family expansions in various groups that are targets for novel drug and vaccine development. Additional more focused comparative genomic work has revealed differences between helminth and host metabolic pathways and extracellular vesicle protein content that represent potential druggable targets, evidence for positive selection in gene families that are uniquely expanded in flatworms and implicated in the biology of endoparasitism, conserved immunomodulatory proteins that are potential vaccine targets for soil-transmitted helminths, and patterns of coevolution between parasitic roundworms and their mammalian hosts^8–14^. These advances further highlight helminth reference genomes as indispensable tools for comparative biological and biomedical research.

Reductions in the cost and improved ease of access of Next-Generation Sequencing (NGS) has made the assembly of whole genomes from helminths more feasible^15^. As a result, reference genomes have now been generated for the majority of species of medical and veterinary importance^16, 17^. Most of these species, however, are still represented by only a single reference genome. These solitary references are often generated from laboratory models that have been maintained for decades and therefore cannot represent the biological diversity observed in natural helminth populations^18, 19^. Generating multiple reference genomes per species from contemporary and geographically disparate populations would begin to capture the genomic diversity of helminths, and allow for characterization of differences in, for example, genome organization, gene copy number and structural variants, and adaptation to local selective pressures. When generating a genome from a natural population, using genomic DNA from one specimen is ideal to avoid complications in the assembly process. Therefore, streamlined laboratory and computational workflows that allow high-quality assemblies to be generated from an individual specimen using a single data source would be invaluable. The Oxford Nanopore Technologies (ONT) MinION is a single molecule sequencing platform capable of generating long reads ideal for *de novo* genome assembly. Lower read-level accuracy of ONT data has previously required long-read assemblies to be error-corrected with more accurate short-read data. Recent updates to ONT chemistries, however, have decreased the error rate of MinION reads, allowing for the potential to generate helminth reference genomes using only long-read data.

This study aims to assess the contiguity, completeness, gene content, and gene composition of whole genome *de novo* assemblies of parasitic nematodes using MinION data only. Three species that collectively span the breadth of parasitic nematode diversity were utilized for sequencing: the filarial worm *Brugia malayi* (Spirurina), the whipworm *Trichuris trichiura* (Trichinellida), and the dog hookworm *Ancylostoma caninum* (Rhabditina). Assemblies generated using only ONT MinION data were compared to reference genomes for each species, as well as to assemblies supplemented with short-read Illumina data through polishing or hybrid assembly approaches that were generated as a part of this study. This work highlights a straightforward approach for generating high-quality *de novo* genome assemblies from parasitic nematodes using only data generated from the ONT MinION.

## Results

### Data generation

Total gDNA was successfully extracted from a single adult worm for each of *B. malayi* and *T. trichiura*, and from a pool of L3 larvae for *A. caninum*. For each species, both MinION and Illumina sequencing libraries were prepared from the same source of gDNA. The bioinformatic pipeline used to generate each assembliy type is outlined in Figure 1. Total amounts sequence data generated for each species prior to basecalling and/or quality control were as follows: ∼15.9 Gb (MinION) and ∼7.9 Gb (Illumina) for *B. malayi*; ∼26.8 Gb (MinION) and ∼4.5 Gb (Illumina) for *T. trichiura*; and ∼33.1 Gb (MinION) and ∼20.8 Gb (Illumina) for *A. caninum* (Supplemental Table 1). For all three species, a median quality score value of >Q16 was achieved for MinION reads, and for MinION reads aligned to final MinION data only assemblies (Supplemental Figure 1). Values for average depth of coverage of quality-controlled reads mapped to species-specific reference assemblies were as follows: 124.85× (MinION) and 82.95× (Illumina) for *B. malayi*; 249.91× (MinION) and 48.58× (Illumina) for for *T. trichiura*; and 49.42× (MinION) 38.64× (Illumina) for *A. caninum*. For all species and assembly types, complete mitochondrial genomes were concurrently generated. For *B. malayi*, the complete genome for the *Wolbachia* endosymbiont was also assembled in a single circular genome.

**Figure 1.**
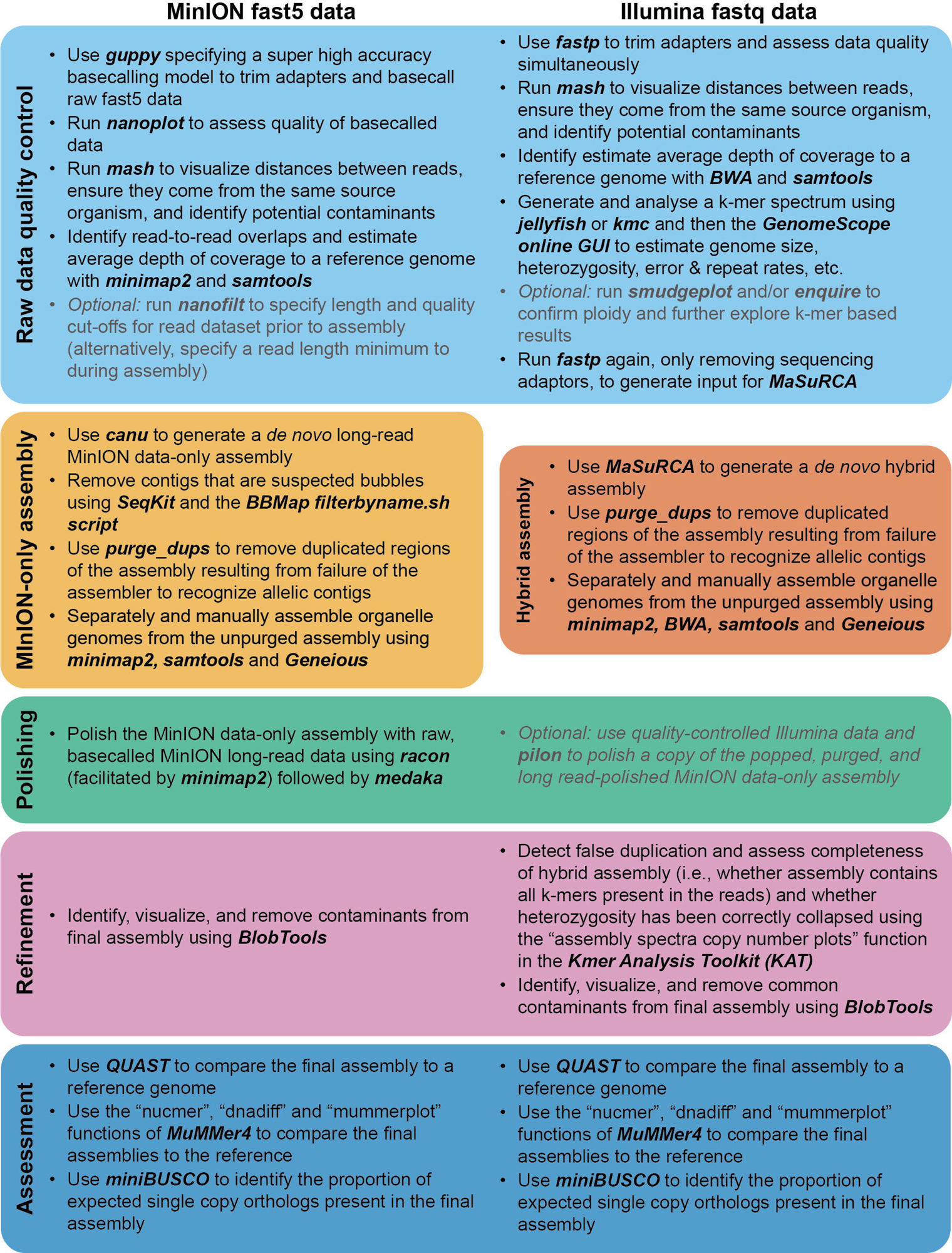
Bioinformatic pipelines used to generate *de novo* whole genome assemblies. The MinION data only pipeline is provided at left and the hybrid pipeline is provided at right.

### Assembly contiguity

All assemblies generated in this study were shorter in terms of total length compared to current reference genomes, with the exception of the MinION data only assembly for *B. malayi* (Table 1). Of the assemblies generated in this study, hybrid assemblies were consistently the most contiguous as represented by higher N50 values (Table 1). The most improvement in terms of contiguity was observed in *A. caninum*, where each assembly approach resulted in a shorter genome and more contiguous assembly compared to the current reference genome. Within a species, all assemblies had similar levels of GC content.

**Table 1.** Comparative quality metrics output by QUAST for the assemblies generated as part of this study and the reference assemblies available for each species. Abbreviations: bp=base pairs.

|  |  | **Length (bp)** | **No. contigs** | **N50 (bp)** | **GC content** | **N content** |
| --- | --- | --- | --- | --- | --- | --- |
| *Brugia malayi* | **Ghedin et al. (2007) reference assembly** | 88,235,797 | 197 | 14,214,749 | 28.5% | 0.30% |
|  | **MinION data only assembly** | 88,513,498 | 88 | 4,666,641 | 28.43% | 0% |
|  | **MinION assembly polished with Illumina data** | 88,428,546 | 88 | 4,667,161 | 28.45% | 0% |
|  | **Hybrid assembly** | 84,528,932 | 49 | 3,943,761 | 28.57% | <0.001% |
| *Trichuris trichiura* | **Foth et al. (2014) reference assembly** | 75,474,068 | 4,086 | 70,602 | 42.3% | 0.30% |
|  | **Doyle et al. (2022) reference assembly** | 80,573,711 | 113 | 11,299,416 | 42.3% | 0.31% |
|  | **MinION data only assembly** | 73,094,541 | 174 | 806,679 | 42.22% | 0% |
|  | **MinION assembly polished with Illumina data** | 72,959,001 | 174 | 805,117 | 42.23% | 0% |
|  | **Hybrid assembly** | 72,705,678 | 63 | 2,773,302 | 41.15% | 0% |
| *Ancylostoma caninum* | **International Helminth Genomes Consortium (2019) reference assembly** | 465,750,606 | 25,339 | 256,700 | 42.8% | 13.50% |
|  | **MinION data only assembly** | 358,159,610 | 1,578 | 638,901 | 43.36% | 0% |
|  | **MinION assembly polished with Illumina data** | 357,986,713 | 1,578 | 637,973 | 43.38% | 0% |
|  | **Hybrid assembly** | 348,034,555 | 652 | 1,095,227 | 43.24% | <0.001% |

### Assembly completeness

miniBUSCO scores were nearly identical among the three assembly types generated for all three species and matched or exceeded completeness scores of current reference assemblies (Table 2). For *A. caninum*, miniBUSCO scores indicate that the assemblies generated here are more complete than the existing International Helminth Genomes Consortium^7^ reference assembly. All assemblies compared for *T. trichiura*, including both available references, had high proportions of missing BUSCOs (i.e., ∼40.8–42.4%; see Table 2). According to the assessment of contiguity versus completeness, the assemblies generated here for *B. malayi* and *A. caninum* can be classified as “tier 1” genomes (Figure 2). For *T. trichiura*, the contiguity of assemblies generated here is sufficient to qualify them each as “tier 1”, but, as mentioned above, high proportions of BUSCO missingness prevented any trichurid assembly assessed from achieving “tier 1” status (see Figure 2).

**Figure 2.**
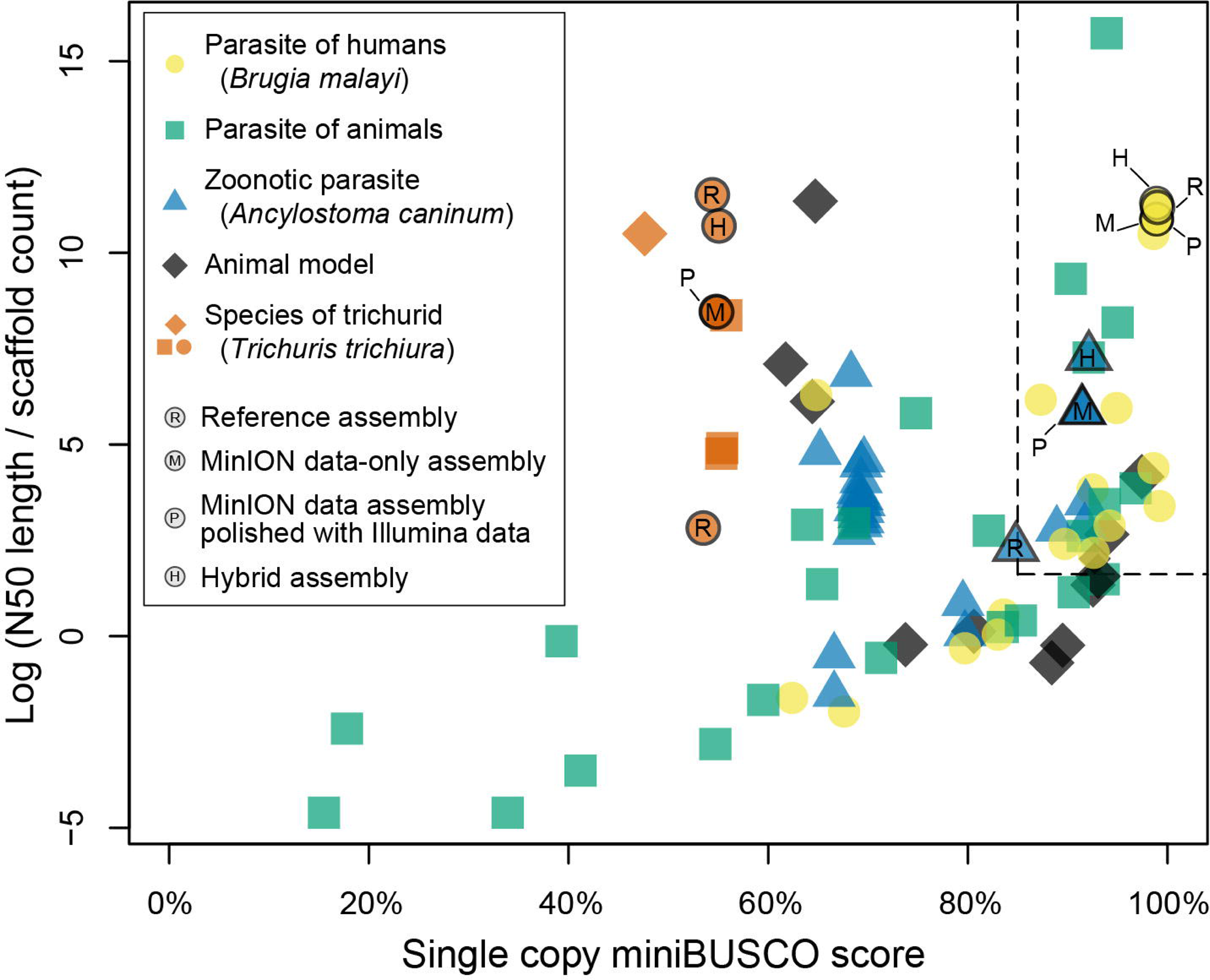
Plot of single copy miniBUSCO score versus the log value of N50 length divided by contig (or scaffold) count for the assemblies generated as part of this study, and for all genomes of nematodes that parasitize animals as adults available from WormBase ParaSite. Dotted lines indicate the cutoff for “tier 1” genome status *sensu* the International Helminth Genomes Consortium (i.e., >85% single copy BUSCO score and >1.6 log value for the contiguity metric). A key to symbolic designations is provided at top left.

**Table 2.** Scores from miniBUSCO for the assemblies generated as part of this study and the reference assemblies available for each species. Scores are presented as number of BUSCOs recovered in each assembly followed in parentheses by proportion of the total number of nematode orthologs assessed by miniBUSCO (n=3,131).

|  |  | Single copy | Duplicated | Fragmented | Missing |
| --- | --- | --- | --- | --- | --- |
| *Brugia malayi* | Ghedin et al. (2007) reference assembly | 3,100 (99.01%) | 15 (0.48%) | 8 (0.26%) | 8 (0.26%) |
|  | MinION data only assembly | 3,097 (98.91%) | 17 (0.54%) | 9 (0.29%) | 8 (0.26%) |
|  | MinION assembly polished with Illumina data | 3,097 (98.91%) | 17 (0.54%) | 9 (0.29%) | 8 (0.26%) |
|  | Hybrid assembly | 3,096 (98.88%) | 17 (0.54%) | 8 (0.26%) | 10 (0.32%) |
| *Trichuris trichiura* | Foth et al. (2014) reference assembly | 1,676 (53.53%) | 15 (0.48%) | 112 (3.58%) | 1,328 (42.41%) |
|  | Doyle et al. (2022) reference assembly | 1,702 (54.36%) | 46 (1.47%) | 102 (3.26%) | 1,281 (40.91%) |
|  | MinION data only assembly | 1,717 (54.84%) | 20 (0.64%) | 99 (3.16%) | 1,295 (41.36%) |
|  | MinION assembly polished with Illumina data | 1,716 (54.81%) | 20 (0.64%) | 100 (3.19%) | 1,295 (41.36%) |
|  | Hybrid assembly | 1,725 (55.09%) | 22 (0.70%) | 104 (3.32%) | 1,280 (40.88%) |
| *Ancylostoma caninum* | International Helminth Genomes Consortium (2019) reference assembly | 2,654 (84.77%) | 239 (7.63%) | 143 (4.57%) | 95 (3.03%) |
|  | MinION data only assembly | 2,865 (91.50%) | 100 (3.19%) | 66 (2.11%) | 100 (3.19%) |
|  | MinION assembly polished with Illumina data | 2,867 (91.57%) | 99 (3.16%) | 65 (0.08%) | 100 (3.19%) |
|  | Hybrid assembly | 2,883 (92.08%) | 111 (3.55%) | 57 (1.82%) | 80 (2.56%) |

### Gene content and composition

For organelle genomes, within a species, the mitochondrial and *Wolbachia* genomes generated were nearly identical (i.e., >99.9% pairwise nucleotide identity) across assembly types. For gene datasets produced by GeMoMa, the number of predicted genes was roughly equal across assembly types for each species (Figure 3). Additionally, the majority of these genes were shared between assembly types. Within a species, genes predicted across assembly types were similar or identical in mean gene length, mean length of introns, exons, and coding sequences, and mean number of exons per coding sequence (Table 3). Pairwise nucleotide comparisons of genes shared between MinION only data assemblies and MinION assemblies polished with Illumina data showed 88–98% of these genes to be identical at the nucleotide level (Figure 4A). The majority of differences in gene composition were the result of single SNPs or single indels, with few shared genes demonstrating greater than ten mismatches (Figure 4B).

**Figure 3.**
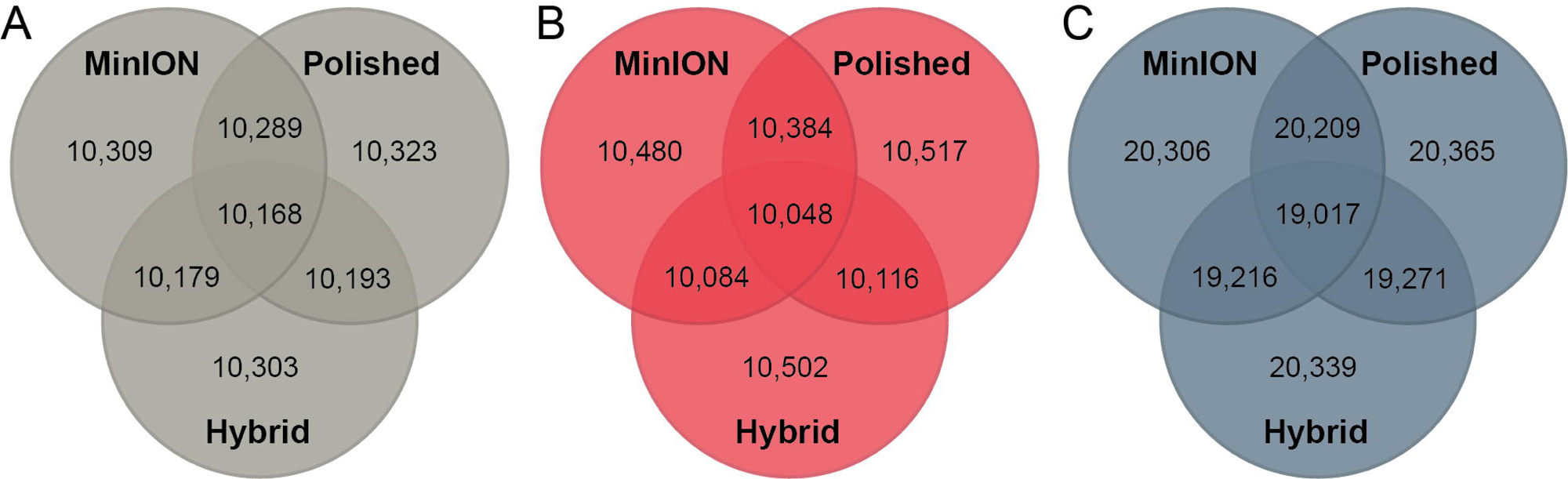
Venn diagrams of the number of genes predicted by GeMoMa that were shared among the assembly types generated. (A) *Brugia malayi*, (B) *Trichuris trichiura* and (C) *Ancylostoma caninum*.

**Figure 4.**
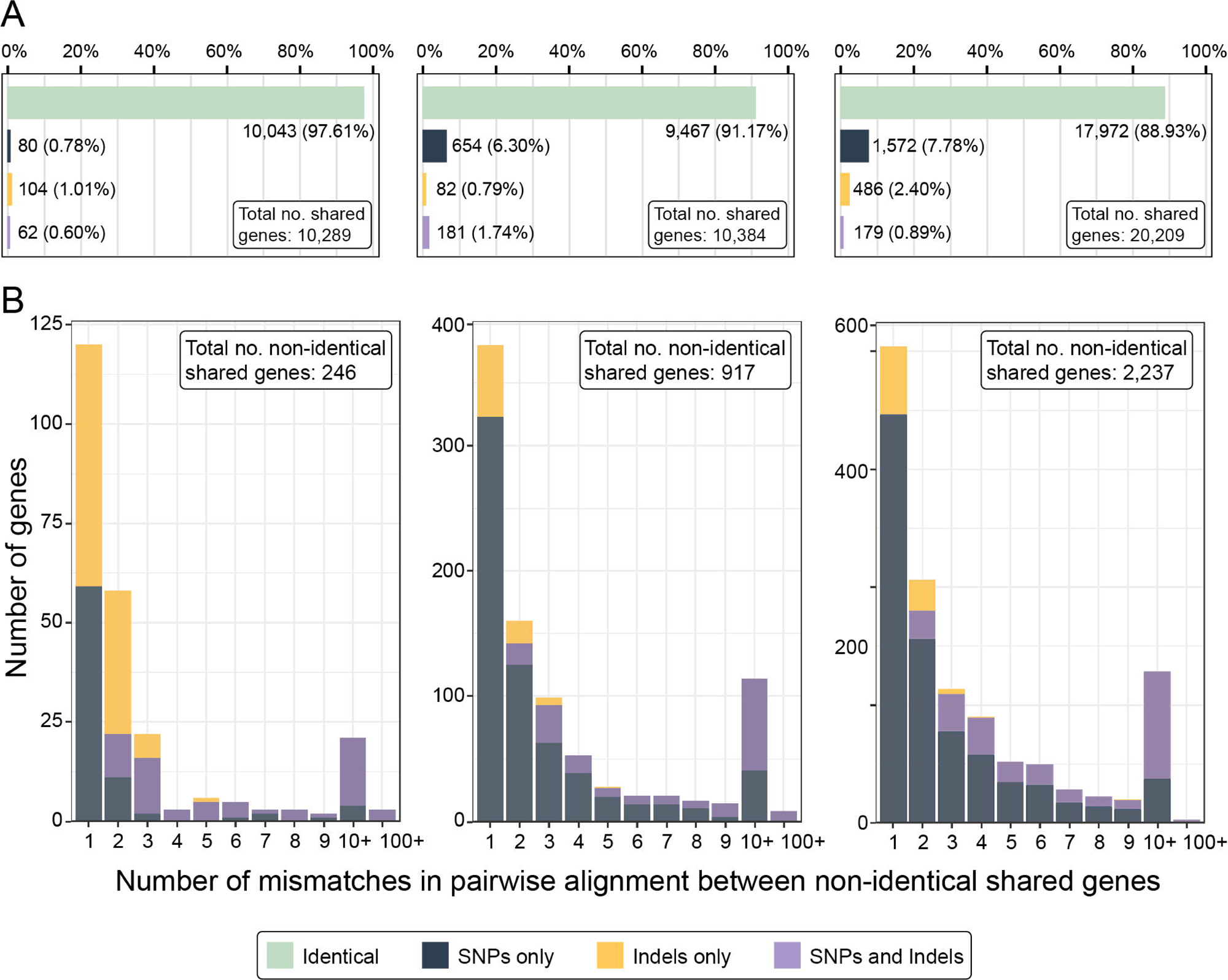
Summary of differences in gene composition identified in predicted genes shared between MinION data assemblies and MinION assemblies polished with Illumina data. **(A)** Proportions of shared genes identified at the nucleotide level as identical, differing by SNPs only, differing by indels only, or differing by both SNPS and indels for *Brugia malayi* (left), *Trichuris trichiura* (center), and *Ancylostoma caninum* (right). **(B)** Number of differences identified between non-identical shared genes for *Brugia malayi* (left), *Trichuris trichiura* (center), and *Ancylostoma caninum* (right). A key to bar plot colors is provided at bottom center.

**Table 3.**
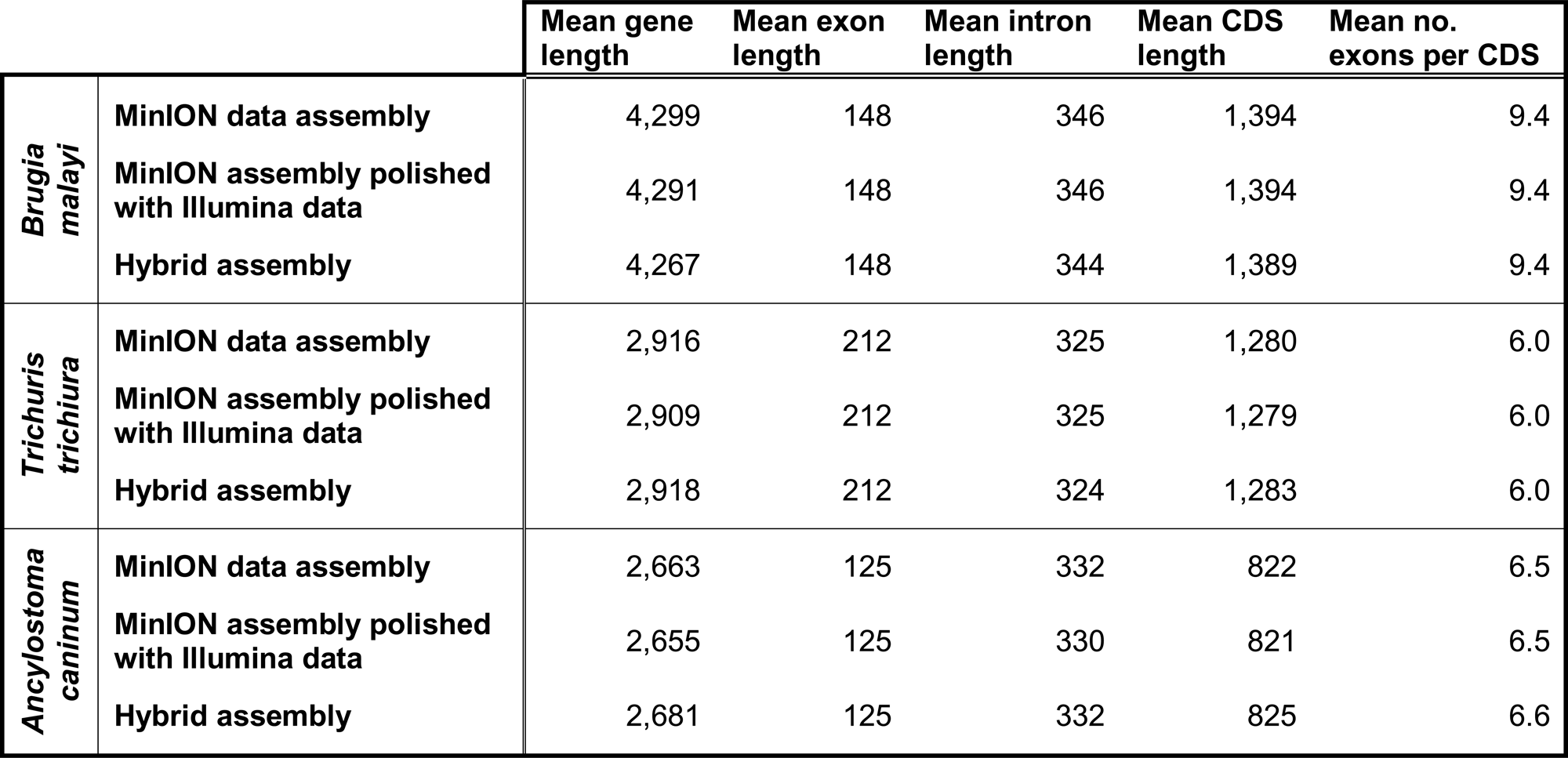
Comparative summary statistics output by AGAT for genes predicted by GeMoMa in assembly types for each species. Mean lengths are presented as number of base pairs. Abbreviations: CDS=coding sequence.

## Discussion

### Sample processing and sequencing

The DNA extraction and MinION library preparation protocols described here were optimized for low input gDNA extracted from individual parasitic nematodes, allowing us to retain the majority of input gDNA through library preparation. Genomic DNA from a single individual is the ideal input for *de novo* whole genome assembly for diploid organisms. This allows assembly pipelines to contend with only two potential haplotypes, leading to more accurate assemblies with less haplotypic duplication. An alternative method would be to sequence multiple individuals from a highly inbred homozygous line. Many species cannot be maintained in a laboratory setting, however, making the establishment of inbred lines for these taxa challenging or impossible^20^. Additionally, sampling from natural, rather than laboratory-maintained, populations is preferred for many genomic studies, further highlighting the advantage of whole-genome sequencing from single individuals^21^. For *B. malayi* and *T. trichiura*, which both have relatively large adult stages (∼3–5 cm in length) and tractable genome sizes (<100 Mb), optimized protocols allowed for the generation of both short- and long-read data from a single adult worm. Adults of *A. caninum*, however, tend not to exceed ∼1.5 cm in length, and have the largest genome size of the three focal species in this study (i.e., ∼400 Mb). Thus, for *A. caninum*, even if adult worms had been accessible, these protocols would likely still not have allowed for generation of sufficient genomic data from a single adult. Despite using a pooled sample, however, copy number spectrum plots suggest that incorporation of a purging step produces results similar to those for single individuals (see Supplemental Figure 2). For nematodes where gDNA from single individuals does not meet the requisite quantity for sequencing on the ONT MinION, additional strategies such as whole genome amplification can be pursued.

There was an association between the average Q score of MinION read datasets and the total amount of data generated by a flow cell: More efficient MinION sequencing runs generated data with higher average Q scores (see Supplemental Table 1; Supplemental Figure 1). For example, MinION libraries for both *B. malayi* and *T. trichiura* generated ∼10.8–11.8 Gb of data per flow cell post-basecalling with comparably high Q scores. Conversely, libraries for *A. caninum* generated ∼7.7–8.5 Gb of data per flow cell post-basecalling, and these data had the lowest average Q score of the three species sequenced (see Supplemental Table 1; Supplemental Figure 1). We observed that flow cells that were used to sequence the *A. caninum* libraries experienced the most rapid decline in pore availability. This observation highlights the variability in data generation and quality depending on the sample type when using the latest ONT chemistries.

### Estimation of genome size and heterozygosity

There were appreciable differences in overall size of assemblies produced in this study compared to reference genomes (see Table 1). These differences were notable for both *T. trichiura* and *A. caninum*, but most dramatic for *A. caninum*, where assemblies produced here were >100 Mb smaller than the existing reference. This is likely due to the fact that a purging step to remove haplotypic duplication was included in the bioinformatic pipeline for all assembly types generated in this study (see Figure 1). This was not the case for the reference genome for *A. caninum*^7^, and thus it likely includes haplotypic duplication that artificially inflates genome size. Three lines of evidence support that the genomes assembled here more accurately represent the true haplotypic genome size for both *T. trichiura* and *A. caninum*. First, the lengths of these assemblies more closely match k-mer based genome size estimates than do those of the existing reference genomes (see Table 1). Second, miniBUSCO scores were largely indistinguishable across all assembly types (including existing reference assemblies), indicating that all are comparably complete despite ranging in size. In fact, the Doyle et al. reference assembly for *T. trichiura*^22^ and the International Helminth Genomes Consortium reference assembly for *A. caninum*^7^ had higher proportions of duplicated BUSCOs as compared to the assemblies generated herein (see Table 2). Finally, for hybrid assemblies, copy number spectrum plots suggest they were not “over-purged” and thus missing genomic content (see Supplemental Figure 2). As previously advocated by other authors, purging haplotypic duplication is an important step in genome curation regardless of assembly approach^23, 24^. Unsurprisingly, purging becomes increasingly important for highly heterozygous sample types, as observed here for *A. caninum*.

Results from GenomeScope (see Supplemental Table 1) and copy number spectrum plots (see Supplemental Figure 2) indicate low heterozygosity for *B. malayi* and elevated heterozygosity for *T. trichiura* and *A. caninum*. These results are congruent with the fact that the single individual of *B. malayi* sequenced came from a long-maintained inbred laboratory strain while the individual of *T. trichiura* sequenced came from a naturally infected human host, and sequence data for *A. caninum* were generated from a pool of individuals. When sequencing adult females that are potentially gravid, as was the case for *B. malayi*, analyses like GenomeScope and KAT that utilize k-mer spectra can be altered by the presence of paternal haplotypes in eggs. Given that the female of *B. malayi* sequenced came from an inbred strain, however, maternal and paternal haplotypes are likely to be identical, or highly similar, ameliorating this concern. It is worth noting that the k-mer based approaches used to estimate genome size and heterozygosity and to assess the effectiveness of purging rely on short-read data as input, and therefore cannot be used to evaluate long-read only assemblies at this time.

### Assembly contiguity

The reference genome assemblies for both *B. malayi* and *T. trichuris* are highly contiguous and nearing chromosome scale. For *B. malayi* and *T. trichuris*, the N50 values for the assemblies generated in this study were substantially lower compared to those of the highly contiguous reference genome. This is due in part to the additional data sources used to generate these references, which for *B. malayi* included optical mapping^25^ and for *T. trichiura* included unspecified updates that resulted in improvements to contiguity^22^. For *A. caninum*, however, the contiguity of the assemblies generated here represent marked improvements as compared to the reference genome. Hybrid assemblies were in all cases more contiguous than MinION data assemblies (see Table 1). This result was somewhat unexpected given that including short-read data in whole genome assembly has traditionally been touted to increase accuracy rather than contiguity^26–28^. Increased contiguity of hybrid assemblies over long read-only assemblies has been observed in bacteria whole genome sequencing^29, 30^ but does not appear to have been reported previously for eukaryotes. It is worth noting that the long reads generated for each species here did not approach the read lengths that MinION devices are capable of sequencing. Across species, read length N50 values ranged from ∼2.7–8.6 kb (see Supplemental Table 1), which are relatively small. This may be due to the optimized MinION library preparation approach used here, which relied on washing final libraries with a titration of Short and Long Fragment Buffer to retain some short fragments of gDNA in addition to long fragments (see Methods). Rigorous size selection which could have improved read lengths was therefore sacrificed to enable sequencing from low quantities of input gDNA. Had it been possible, generation of ultra-long reads would have likely lessened the gap in contiguity metrics between MinION data and hybrid assemblies. Additionally, a single assembly algorithm was chosen to generate long-read and hybrid assemblies (i.e., Canu and MaSuRCA, respectively). Given the same read-level data, different assembly algorithms may have produced assemblies with different contiguity metrics.

### Assembly completeness

In general, completeness values as measured by miniBUSCO are indistinguishable between assembly types for *B. malayi* and *T. trichiura*. Assemblies generated here for *A. caninum* showed an improvement in the number of single copy genes identified and a reduction in the number of duplicated and fragmented genes identified as compared to the reference genome. Results from miniBUSCO indicate a high proportion of missingness for all trichurid assemblies (see Table 2; Figure 2). Underrepresentation in the BUSCO “nematoda_odb10” dataset has been noted to result in biases and underestimation in ortholog detection for some species of nematodes^31^. This presents a challenge for using traditional completeness metrics like miniBUSCO to assess trichurid genome assemblies.

### Gene content and composition

In terms of gene content, within a species, similar numbers of genes were predicted by GeMoMA for each of the three assembly types generated (see Figure 3). Though not all predicted genes were identified in all three assemblies, large proportions were found to be shared among them (i.e., ∼93–98%, depending on the species). In particular, MinION only assemblies and MinION assemblies polished with Illumina data were highly similar in terms of gene content, with ∼98.7–99.7% of predicted genes shared between them within a species. Furthermore, predicted genes sets had, by species, identical or nearly identical mean genes lengths, mean intron, exon, and coding sequence lengths, and mean numbers of exons per coding sequence (see Table 3), providing further evidence for a high degree of similarity in gene content across assembly types.

In terms of gene composition, a major goal of this study was to assess the potential of Illumina data to correct errors that may be present in the final assembly as a result of less accurate MinION data. To evaluate this, gene datasets predicted by GeMoMa for MinION data only assemblies and for MinION assemblies polished with Illumina data were pairwise aligned and assessed for mismatches. Comparisons between MinION data only assemblies and hybrid assemblies are not tenable because long-read assemblers like Canu and hybrid assemblers like MaSuRCA utilize assembly algorithms that handle read-level data dissimilarly^32, 33^. Given that gene predictions for each assembly were based on existing reference genomes and their corresponding annotations (see Methods), plus these reference genomes were generated from different biological samples, comparison to reference genomes was also not salient.

The most direct comparison to determine whether short reads improve gene accuracy in final assemblies is the MinION data only assembly versus that same assembly polished with Illumina short reads. If Illumina polishing substantially improves accuracy, homology-based gene prediction would be expected to result in a low proportion of genes shared between these two assembly types, and or/the majority of shared genes to differ at the nucleotide level. This was not the case as all three species showed high proportions of predicted genes shared between assembly types, and the majority of the genes shared between the two assembly types were identical (see Figures 3, 4A). For those that were not identical, most differed by only a single SNP or indel. These single differences are presumably corrections made during Illumina polishing (Figure 4B). Despite this, a substantial number of gene comparisons still contained 10–100+ mismatches (Figure 4B). It is unlikely these differences result from polishing errors in MinION data only assemblies; rather, they are more likely the result of homology-based gene prediction models identifying incongruent genes between assembly types in a small number of cases. In summary, depending on the species, correcting with short reads did not change nucleotide calls in coding sequences for ∼88–98% of the genes compared.

## Conclusions

For this study, *de novo* whole genome assemblies were generated for three species of parasitic nematodes (*Brugia malayi*, *Trichuris trichiura*, and *Ancylostoma caninum*) using only MinION long-read data, using MinION data polished with Illumina short reads, and using a combined hybrid approach. For *B. malayi* and *T. trichiura*, optimized gDNA extraction and library preparation protocols allowed for the generation of complete genomes from individual adult worms. For all species sequenced, MinION data only assemblies had similar, or superior, measures of contiguity and completeness as compared to existing reference genomes. The most substantial improvements in quality metrics were observed for *A. caninum*, which was the only one of the three focal species for which the existing reference genome is not a near-chromosome scale assembly. Among the three assembly types generated, predicted gene content was nearly identical with a species, and the vast majority of predicted genes shared between MinION data only assemblies and MinION assemblies polished with short reads were identical at the nucleotide level. For some genes, however, polishing did result in the correction of single SNPs or indels. Although additional data types beyond MinION long reads are needed to produce reference-quality, chromosome-scale assemblies, the results of this study demonstrate that MinION data alone can be used to generate highly contiguous and complete *de novo* whole genomes from parasitic helminths.

## Limitations of the Study

This study has multiple limitations. For two of the three focal species, gDNA was generated from adult worms. For most species, and particularly those that infect humans, adult nematodes are difficult or impossible to obtain, somewhat limiting the utility of this approach in practice. Furthermore, the assemblies generated here remain incomplete in terms of the modern reference genome standards. They are not chromosome-scale, they are not annotated, and additional data types are required to complete them. Generating chromosome-scale annotated assemblies was not the goal of this study. Rather, it was to evaluate the feasibility and quality of whole genome assemblies generated from an accessible NGS data type. For *B. malayi* and *T. trichiura*, existing reference genomes were already exceptionally high-quality “tier 1”-status assemblies *sensu* the International Helminth Genomes Consortium^7^, and the assemblies generated here using a single data source approached their quality in contiguity and matched their quality in completeness. The comparison in which the added value of long-read MinION data is most evident is that of *A. caninum*. Using gDNA from the same strain and sample type on which the existing reference assembly was based, the genomic resources available for this species were improved considerably.

## Author Contributions

KSH and JRF designed the study. JMH provided the specimens of *Ancylostoma caninum* sequenced. PN provided the specimen of *Trichuris trichiura* sequenced. KSH extracted genomic DNA and prepared and sequenced MinION libraries. KSH and JRF determined and refined bioinformatic pipelines. KSH performed bioinformatic analyses. RW aided KSH in gene-level comparison analyses. KSH and JRF interpreted data and generated figures and tables. KSH and JRF drafted the manuscript. All authors contributed to revising and approving the final version of the manuscript.

## Acknowledgements

The authors thank the BEI Resources NIH/NIAID Filariasis Research Reagent Resource Center for providing specimens of *Brugia malayi* for sequencing. Thanks are also due to Dr. Michael Wiley, Dr. Shaun Cross, Kristen Bernhard, and Mahmood Al-Haehm (University of Nebraska Medical Center) for providing access and support for use of select laboratory equipment. The authors additionally thank Christopher Castaldi and Irina Tikhonova (Yale Center for Genome Analysis, Yale School of Medicine) for generation of, and correspondence regarding, Illumina sequence data. Bioinformatic analysis was enabled by the University of Nebraska-Lincoln Holland Computing Center computing cluster.

## Materials and Methods

### Specimen acquisition

The specimens sequenced were obtained from the same strains previously used to generate existing reference genomes for each species. Adults of *B. malayi* (FR3 strain) were acquired through the NIH/NIAID Filariasis Research Reagent Resource Center via BEI Resources (Michalski et al., 2011). Worms were received frozen and stored at -80°C. Prior to sequencing, worms were thawed at 4°C and subsequently preserved in DNA/RNA Shield (Zymo Research, Irvine, CA, USA). Adults of *T. trichiura* were obtained from a Danish patient after anthelmintic treatment for an infection acquired in Uganda. Worms were received preserved in 70% ethanol and were stored at 4°C prior to sequencing. Pooled third-stage (L3) larvae of *A. caninum* (Baltimore strain) were obtained from experimental infection in laboratory-maintained canines. Pooled L3s were received frozen, stored at -80°C, and thawed at 4°C prior to sequencing.

### DNA extraction

Total nucleic acid was extracted separately from each of a single adult female (*B. malayi*), a single adult male (*T. trichuris*), and a pool of L3 larvae (*A. caninum*) using a *Quick*-DNAÔ HMW MagBead Kit (Zymo Research, Irvine, CA, USA) and modified Solid Tissue extraction protocol (Supplemental Table 1). Prior to extraction, adult worms were placed in clean 1.5 mL DNA LoBind® tubes (Eppendorf® North America, Enfield, CT, USA) using sterilized forceps, while pooled L3s were centrifuged at 6,540 *g* for 3 min to allow for the removal of excess supernatant. All samples were then homogenized mechanically using a Monarch® Pestle Set single-use microtube pestle (New England Biolabs® Inc., Ipswich, MA, USA). Homogenization was performed both prior to, and immediately following, the addition of Proteinase K. Samples were then digested in a digital dry bath at 55°C for 24 hr with occasional flick mixing. The protocol for DNA purification was then followed, with these modifications: After the addition of 33 µL of MagBinding Beads, samples were placed on a benchtop rotating mixer for 40 min to 1.5 hr at RT; and after the addition of 52 µL DNA Elution Buffer, samples were incubated in a digital dry bath at 37°C for 2 hr with occasional flick mixing, then incubated at RT overnight prior to collecting eluted DNA. To avoid DNA fragmentation, samples were mixed by flicking rather than pipette mixing wherever possible. The concentration of each extraction was measured using a Qubit4Ô 1X dsDNA High Sensitivity (HS) Assay (ThermoFisher Scientific, Waltham, MA, USA) and fragment size distribution was determined using an Agilent 2200 TapeStation System and associated protocol for genomic DNA ScreenTape analysis (Agilent, Santa Clara, CA, USA). For *T. trichiura*, TapeStation analysis was not performed.

For *A. caninum*, initial attempts to sequence ONT libraries resulted in low sequencing efficiency, poor data quality, and rapid pore loss. To ameliorate concerns of protein contamination, extracted genomic DNA (gDNA) was purified via bead-based cleanup prior to final ONT library preparation. This cleanup was performed as follows: Two aliquots of extracted gDNA were each bound to a 0.5× volume of AMPure XP beads (Beckman Coulter, Brea, CA, USA) on a benchtop rotator mixer for 1 hr at RT. Beads were then washed twice with 80% EtOH, air-dried, and gDNA was eluted in 52 µL Zymo DNA Elution Buffer for 2 hr at 37°C with occasional flick mixing, followed by overnight elution at RT. This cleanup process was repeated a second time for both aliquots. Purified extractions were then quantified using a Qubit4^TM^ 1X dsDNA High HS Assay and the Agilent 2200 TapeStation System.

### MinION library preparation

Aliquots of gDNA extracted for each species were used to prepare one or more libraries for whole genome sequencing on an ONT MinION desktop sequencer (see Supplemental Table 1). Libraries were prepared using a combination of the DNA Clean & Concentrator MagBead (Zymo Research, Irvine, CA, USA) and ONT SQK-LSK114 Ligation Sequencing (Oxford Nanopore Technologies, Oxford, United Kingdom) Kits, and a modified hybrid protocol. First, gDNA was incubated with ONT DNA repair and end preparation reagents for 10 min at RT followed by 10 min at 65°C. End preparation reactions were then bound to 20 µL Zymo MagBinding Beads in 4× volumes of Zymo DNA MagBinding Buffer on a benchtop rotating mixer for 30 min to 2 hr. Beads were then washed twice with Zymo DNA Wash Buffer, air-dried for 10 min, eluted in 51 µL Zymo DNA Elution Buffer via manual agitation for 10 min, and quantified via Qubit 1X dsDNA HS Assay. Sequencing adaptors were then ligated via a 15 min incubation at RT, and libraries were bound to a 0.4× volumes of AMPure XP beads on a benchtop rotator mixer at RT for 1 hr. Beads were then washed twice with either ONT Long Fragment Buffer (*A. caninum*) or a titrated wash mix of 1:3 ONT Short Fragment Buffer:ONT Long Fragment Buffer (*B. malayi* and *T. trichiura*). After washing, beads were air-dried, then libraries were eluted in 15–17 µL of ONT Elution Buffer for 2 hr at 37°C with occasional flick mixing followed by overnight elution at RT. Final libraries were quantified using a Qubit4Ô 1X dsDNA High HS Assay and the Agilent 2200 TapeStation System. For *T. trichiura*, TapeStation analysis was not performed. Additionally, for *T. trichiura*, a portion of ONT library reserved after sequencing on one flow cell was re-washed prior to sequencing on a second flow cell (see Supplemental Table 1). This aliquot of library was bound to a 0.4× volume of AMPure XP beads on a benchtop rotator mixer for 1 hr, washed twice with ONT Long Fragment Buffer, air-dried, then eluted in 17 µL of ONT Elution Buffer for 2 hr at 37°C with occasional flick mixing followed by overnight elution at RT.

### MinION sequencing, basecalling, and read data quality control

Portions of ONT libraries were sequenced for ∼62–80 hr each on one (*B. malayi*), two (*T. trichiura*), or three (*A. caninum*) ONT MinION R10.4.1 flow cells. The amount of library loaded onto each flow cell and the total number of pores available at the start of sequencing are provided in Supplemental Table 1. MinKNOW software (ONT) v. 22.10.7 (*B. malayi*), v. 22.10.10 (*T. trichiura*), or v. 22.12.7 (*A. caninum*) was used to run each flow cell with pore scans every 1.5 hr. After sequencing, signal data (i.e., fast5 files) from each flow cell were basecalled using Guppy v. 6.3.4 (ONT). The bioinformatic pipeline used to generate MinION data assemblies is outlined in Figure 1. Sequencing adaptors were simultaneously removed by specifying the “1717trim_adapters” flag. The results of each Guppy run were summarized using NanoPlot v. 1.40.2^34^. Fastq files that passed basecalling were input to Mash v. 2.2.2^35^ to confirm there was no significant contamination in the read-level data prior to assembly. Estimated genome sizes provided to Mash were based on the lengths of the existing reference assemblies available for each species, and were thus set as 88 Mb for *B. malayi*^36^, 465 Mb for *A. caninum*^7^ and 75 Mb for *T. trichiura*^22^ (https://github.com/stephenrdoyle/ancient_trichuris/tree/master/02_data). Average depth of coverage of reads across the species-specific reference genome and proportion of reads mapped were estimated using Minimap2 v. 2.16^37^ and SAMtools v. 1.9^38^.

### Illumina library preparation, sequencing, and read data quality control

Aliquots of gDNA extracted for each species were sent to the Yale Center for Genome Analysis (YCGA) for Illumina whole genome library preparation and sequencing (see Supplemental Table 1). Libraries were prepared using an xGenTM cfDNA & FFPE DNA Library Prep Kit with unique dual indexing (Integrated DNA Technologies, Newark, NJ, USA) and standard protocol. Libraries were subjected to 5– 7 cycles of PCR to increase concentration (see Supplemental Table 1) and multiplexed for 2×150 paired-end sequencing on an Illumina NovaSeq 6000 targeting 50× depth of coverage genome-wide for each species. Reads were demultiplexed by the YCGA and subsequently filtered for remaining adaptor contamination, quality, and length using fastp^39^. The bioinformatic pipeline used to process Illumina data and generate hybrid assemblies is outlined in Figure 1. Fastq files that passed fastp were input to Mash to confirm there was no significant contamination in the read-level data prior to assembly. Estimated genome sizes provided to Mash were based on the lengths of the existing reference assemblies available for each species (see above). Average depth of coverage across the species-specific reference genome and proportion of reads mapped were estimated using BWA v. 0.7.17 and SAMtools v. 1.9^38^. A second round of fastp was run on the raw demultiplexed Illumina data to remove sequencing adapters, only, by specifying “1717detect_adapter_for_pe”, “1717disable_length_filtering”, and “1717disable_quality_filtering”. These data were the used for hybrid genome assembly, as MaSuRCA utilizes built-in error correction and cleaning.

### Estimation of genome size and heterozygosity

Quality-controlled Ilumina data were used to estimate genome size and heterozygosity for each species. First, Jellyfish v. 2.3.0^40^ was used to generate k-mer spectra, specifying a k-mer size of 21 and an initial hash size of 1,000,000,000. Histograms from Jellyfish were then provided to the GenomeScope web-based graphical user interface^41^ to visualize k-mer spectra and estimate heterozygosity.

### MinION data genome assembly

FASTQ files that passed basecalling by Guppy were concatenated and used as input to Canu v. 2.1^32^ (ONT) to generate MinION data assemblies. For *T. trichiura* and *A. caninum*, a 3 kb minimum read length requirement was specified, and for *B. malayi*, a 5 kb minimum read length requirement was specified. Estimated genome sizes provided to Canu were based on the lengths of the existing reference assemblies available for each species (see above). Contigs in the resulting Canu assemblies that were indicated as potential alternative alleles (i.e., with FASTA headers including “suggestBubble=yes”) were removed using a combination of SeqKit v 0.10.1^42^ and the filterbyname.sh script of BBMap v.38.84^43^. Resulting “popped” assemblies were used as input to purge_dups v. 1.2.5^44^ to remove false duplications. These “popped and purged” assemblies were then polished with one round of Racon v. 1.5.0^45^ followed by one round of Medaka v. 1.7.2 (https://github.com/nanoporetech/medaka) to generate “popped, purged, and polished” assemblies. Copies of these final assemblies were also separately polished with Illumina data using Pilon v. 1.24^46^ and BWA. Three iterations of Pilon were iteratively run specifying the “1717diploid” flag.

### Hybrid genome assembly

FASTQ files that passes basecalling by Guppy and Illumina data from which only sequencing adaptors were removed by fastp were used as input to MaSuRCA v. 4.1.0 (Zimin et al., 2013) to generate hybrid assemblies. Default settings were used except for setting “LHE_COVERAGE” to “35” and “cgwErrorRate=0.15”, and setting “JF_SIZE” to a value ten times the genome size of the species-specific reference assembly. Assemblies output by MaSuRCA were input to purge_dups v. 1.2.5^44^ to remove false duplications.

### Quality control and refinement of final assemblies

To ensure “purged” MaSuRCA hybrid assemblies were not over- or under-purged, copy number spectrum plots were generated using the K-mer Analysis Toolkit (KAT) v. 2.4.0^47^ specifying a k-mer size of 31. “Popped, purged, and polished” Canu MinION data assemblies and “purged” MaSuRCA hybrid assemblies were input to BlobTools v. 1.1.1^48^ to identify potential contaminants. First, the reads used to generate each assembly were mapped to the assembly using Minimap2 (MinION data) and/or BWA (Illumina data), and SAMtools was used to generate sorted BAM files for each mapping. The “blastn” function in BLAST v. 2.14.0^49^ was then used to compare all contigs to reference sequences available in the National Center for Biotechnology Information (NCBI)’s nucleotide databases, specifying “17evalue 1e-25”, “17max_target_seqs 10” and “17max_hsps 1”. BLAST outputs and sorted BAM files were then used as input to BlobTools. Contigs that could be confirmed as host, human, or bacterial contamination were excluded from final assemblies. Quality scores (i.e., Q scores) of basecalled MinION reads, and Q scores of reads when mapped to the final “popped, purged, and polished” MinION data assemblies, were measured using NanoPlot and plotted using the “geom_density” function in the ggplot2 package^50^ in R v. 4.3.0^51^ in RStudio v. 2023.03.1^52^.

### Mitogenome and *Wolbachia* genome assembly

For *B. malayi* and *A. caninum* contigs containing the mitogenomes—and, for *B. malayi*, the genome of its *Wolbachia* endosymbiont—were extracted from “popped” MinION and unpurged MaSuRCA assemblies. For *T. trichiura*, mitogenomes were extracted from the original “unpopped” Canu assembly and the unpurged MaSuRCA assembly. These organelle genomes contigs were then mapped to species-specific reference organelle genomes using Geneious Prime v. 2022.0.1^53^ and manually reconfigured to match the linear orientation of the reference. For *B. malayi* and *T. trichiura*, reference organelle genomes were extracted from the reference genome assemblies for each species (see above). For *A. caninum*, for which a mitochondrial contig is not labeled in the reference assembly, a reference mitogenome was downloaded from GenBank (accession no. MN215971)^54^. Contigs containing complete organelle genomes were then manually added to the appropriate final version of each assembly.

### Assessing assembly contiguity and completeness

Contiguity and completeness were assessed for each assembly using QUAST v. 5.0.2^55^ and miniBUSCO v. 0.2^56^, respectively. For miniBUSCO, the Nematoda ortholog database (nematoda_odb10) was specified. For comparison, miniBUSCO was also run on the aforementioned reference genomes for each species, as well as for all genomes of nematodes that parasitize animals as adults available from WormBase ParaSite^16, 17^. For each assembly, the proportion of single copy BUSCOs present was plotted against the log value of the assembly’s N50 length divided by its scaffold or contig count using R v. 4.2.3 via RStudio v. 2023.06.1. Cutoff values of >85% (for proportion of single copy BUSCOs present) and >1.6 (for the log value of the contiguity metric) were used to assess whether each assembly could be classified as a “tier 1” genome following the standard for helminth genomes established by the International Helminth Genomes Consortium^7^.

### Assessing gene content and composition

Homology-based gene prediction was performed for each assembly generated here using GeMoMA^57, 58^. Predictions were made using default settings and the species-specific reference genome and annotation as input. Resulting GFF files were summarized using AGAT^59^. For each species, the predicted genes shared between the MinION data assembly and the MinION assembly polished with Illumina data were pairwise aligned using the “pairwiseAlignment” function in Biostrings package v. 2.66.0^60^ in R v. 4.2.3 via RStudio v. 2023.06.1 specifying the alignment type as “global”. Biostrings was then used to characterize each pairwise alignment in terms of percent identity, alignment length, gene lengths, and number of single nucleotide polymorphisms (i.e., SNPS) and insertions/deletions (i.e., indels). Summary figures for all comparisons were generated in R v. 4.3.0 via RStudio v. 2023.03.1 using the ggplot2 package.

## Supplemental Figure Titles and Legends

**Supplemental Figure 1.** Histograms of average read quality scores (i.e., Q scores) for unaligned basecalled MinION read data, and for basecalled MinION read data aligned to the final MinION data only genome assembly. (A) *Brugia malayi*, (B) *Trichuris trichiura*, and (C) *Ancylostoma caninum*.

**Supplemental Figure 2.** K-mer multiplicity plots generated from purged hybrid assemblies. (A) *Brugia malayi*, (B) *Trichuris trichiura*, and (C) *Anyclostoma caninum*.

