## Supplemental Table 1 for "Assessing the quality of *de novo* parasite genomes assembled using only Oxford Nanopore Technologies MinION data"

**Supplemental Table 1. Extraction and ONT MinION and Illumina sequence data generation details for each of the focal species sequenced.** Asterisk (*) indicates amount of ONT MinION library remaining after the aliquot of the original library remaining following the first round of sequencing was rewashed with Long Fragment Buffer (LFB).

|  |  | ***Brugia malayi*** | ***Trichuris trichiura*** | | ***Ancylostoma caninum*** | | |
| --- | --- | --- | --- | --- | --- | --- | --- |
| **Extraction** | **Extracted material** | Single adult female | Single adult male | | Pooled L3 larvae | | |
|  | **Total amount of gDNA extracted (ng)** | 737 | 1,508 | | 5,200 | | |
|  | **Mean fragment length of gDNA (bp)** | >60,000 | ––– | | 33,504 | | |
| **Illumina data** | **gDNA input for library prep (ng)** | 42 | 40 | | 400 | | |
|  | **No. PCR cycles** | 7 | 7 | | 5 | | |
|  | **Data pre-quality control and filtering (bp)** | 7,992,851,592 | 4,567,580,276 | | 20,844,376,730 | | |
|  | **Data post-quality control and filtering (bp)** | 7,850,596,985 | 4,442,501,155 | | 20,179,886,997 | | |
| **ONT MinION data** | **Library name** | **Bm ♀ C** | **Tt ♂ 2D (all)** | **Tt ♂ 2D (LFB washed aliquot)** | **Acan L3 B1** | **Acan L3 B2** | **Acan L3 B1 + Acan L3 B2** |
|  | **Mean fragment length of gDNA post-additional bead cleanup (bp)** | ––– | ––– | ––– | 31,955 | 34,563 | ––– |
|  | **Amount of gDNA input for library prep (ng)** | 500 | 1,259 | ––– | 406 | 399 | ––– |
|  | **Amount of library generated (ng)** | 287 | 1,012 | 483* | 189 | 264 | ––– |
|  | **Amount of library sequenced (ng)** | 134 | 78.7 | 76.2 | 164 | 125 | 25 (B1) + 123 (B2) |
|  | **No. pores available at start of sequencing** | 1,413 | 1,544 | 1,635 | 1,527 | 1,465 | 1,538 |
|  | **Total sequencing run time (hr)** | 72 | 72 | 62.75 | 80 | 67.75 | 80 |
|  | **MinKNOW estimated read N50 (kb)** | 8.65 | 2.72 | 5.59 | 6.68 | 6.54 | 6.64 |
|  | **MinKNOW estimated data generated (Gb)** | 15.91 | 14.06 | 12.76 | 10.86 | 10.73 | 11.56 |
|  | **Data post-basecalling with Guppy (bp)** | 11,349,419,698 | 11,891,465,167 | 10,868,016,523 | 8,521,058,505 | 8,261,315,652 | 7,784,337,997 |
| **Assembly** | **Est. depth of coverage for Illumina reads** | 82.95× | 48.58× | | 38.64× | | |
|  | **Proportion of Illumina reads that mapped to reference assembly** | 95.74% | 90.71% | | 90.31% | | |
|  | **Est. depth of coverage for MinION reads** | 124.85× | 249.91× | | 49.42× | | |
|  | **Proportion of MinION reads that mapped to reference assembly** | 98.77% | 99.91% | | 99.85% | | |
|  | **GenomeScope estimated genome size (bp)** | 85,917,606 | 68,538,097 | | 329,957,709 | | |
|  | **GenomeScope estimated heterozygosity** | 0.28% | 1.35% | | 1.09% | | |
