## Supplementary figures and images for "Assessing the quality of *de novo* parasite genomes assembled using only Oxford Nanopore Technologies MinION data"

### Supplemental Figure 1

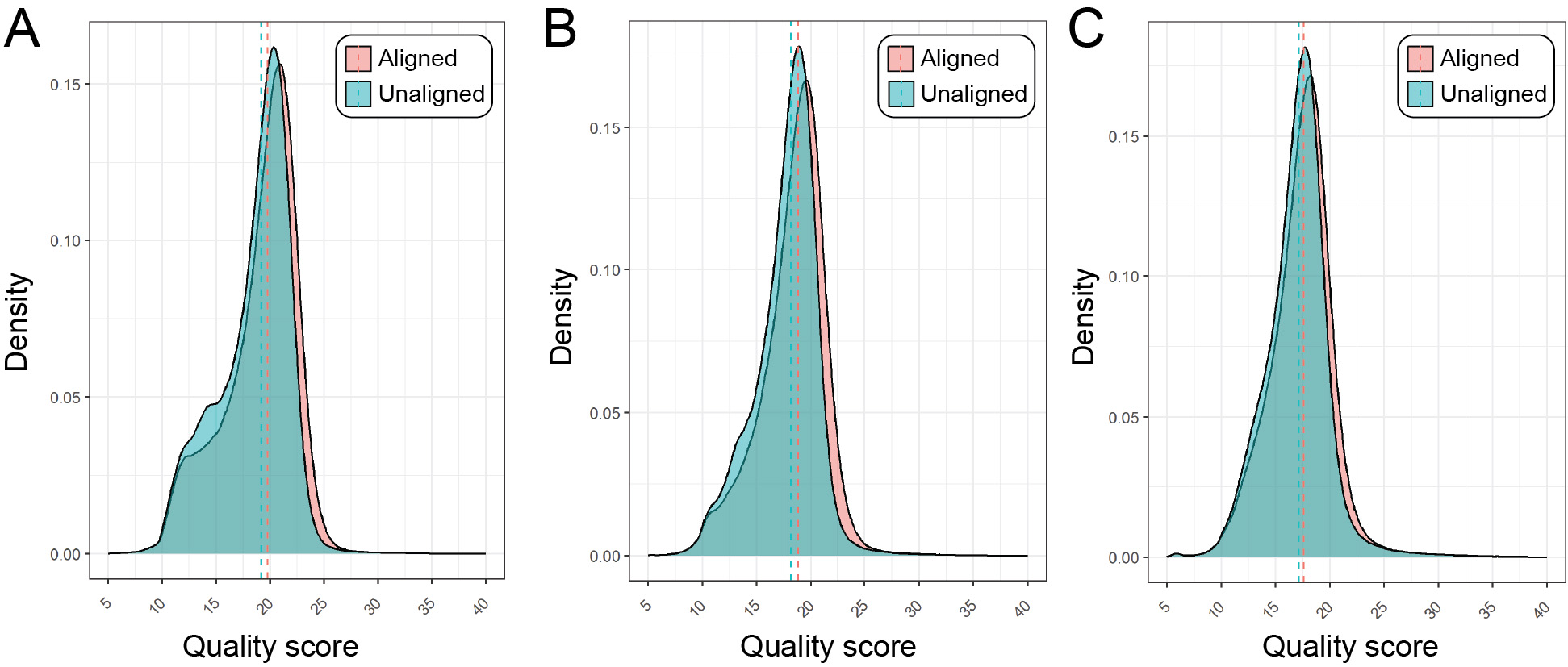

### Supplemental Figure 2

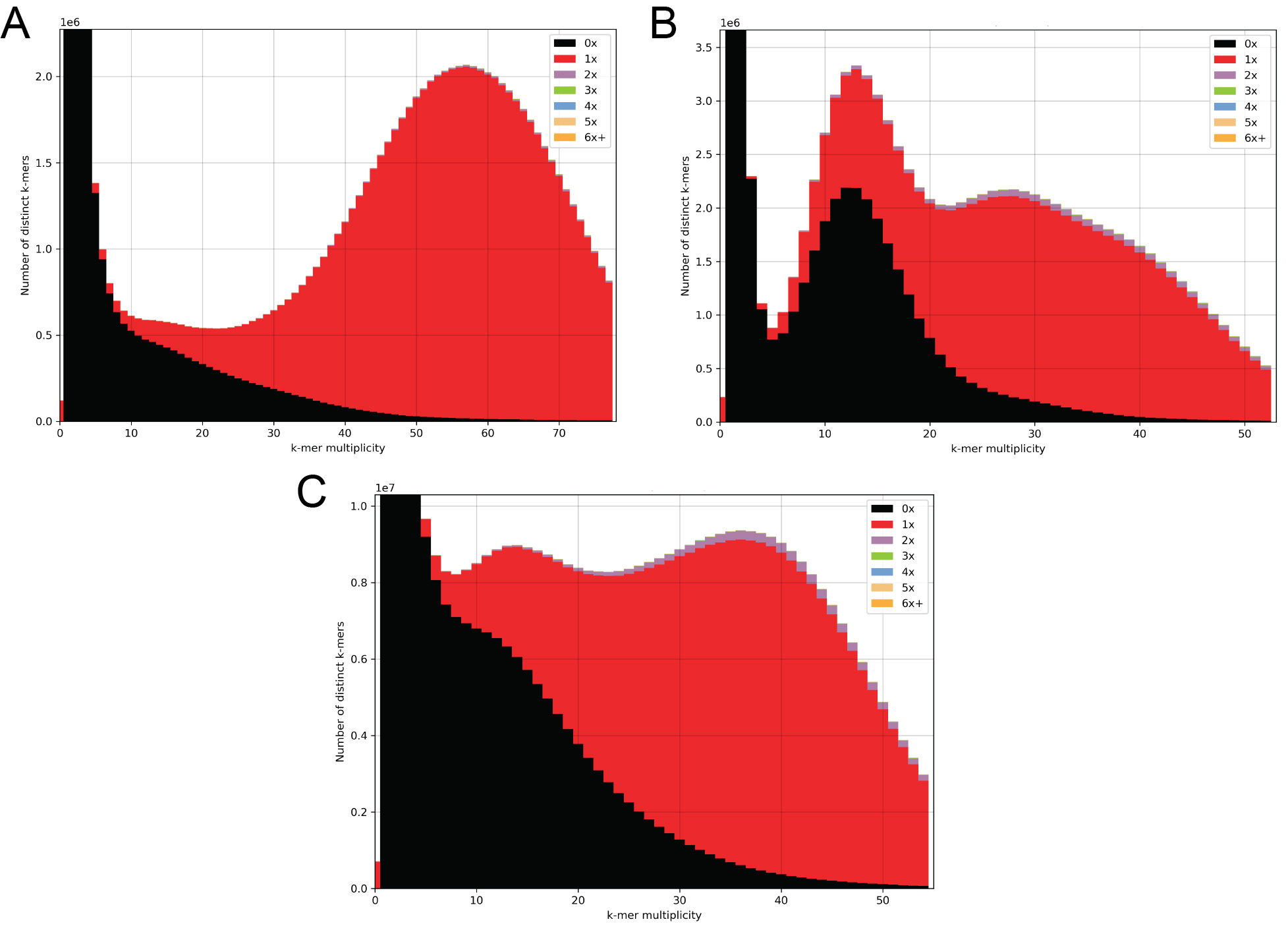
